# Watered-down biodiversity? A comparison of metabarcoding results from DNA extracted from matched water and bulk tissue biomonitoring samples

**DOI:** 10.1101/575928

**Authors:** Mehrdad Hajibabaei, Teresita M. Porter, Chloe V. Robinson, Donald J. Baird, Shadi Shokralla, Michael Wright

## Abstract

Biomonitoring programs have evolved beyond the sole use of morphological identification to determine the composition of invertebrate species assemblages in an array of ecosystems. The application of DNA metabarcoding in freshwater systems for assessing benthic invertebrate communities is now being employed to generate biological information for environmental monitoring and assessment. A possible shift from the extraction of DNA from net-collected bulk benthic samples to its extraction directly from water samples for metabarcoding has generated considerable interest based on the assumption that taxon detectability is comparable when using either method. To test this, we studied paired water and benthos samples from a taxon-rich wetland complex, to investigate differences in the detection of taxa from each sample type. We demonstrate that metabarcoding of DNA extracted directly from water samples is a poor surrogate for DNA extracted from bulk benthic samples, focusing on key bioindicator groups. Our results continue to support the use of bulk benthic samples as a basis for metabarcoding-based biomonitoring, with nearly three times greater total richness in benthic samples compared to water samples. We also demonstrated that few arthropod taxa are shared between collection methods, with a notable lack of key bioindicator EPTO taxa in the water samples. Although species coverage in water could likely be improved through increased sample replication and/or increased sequencing depth, benthic samples remain the most representative, cost-effective method of generating aquatic compositional information via metabarcoding.

---

Aquatic biomonitoring programs are designed to detect and interpret ecological change through analysis of biodiversity in target assemblages such as macroinvertebrates at a given sampling location^1^. The inclusion of biodiversity information in environmental impact assessment and monitoring has injected much-needed ecological relevance into a system dominated by physicochemical data^2^. However, current biomonitoring data suffer from coarse taxonomic resolution, incomplete observation (due to inadequate subsampling), and/or inconsistent observation (variable sampling designs and collection methods) to provide information with sufficient robustness to support the development of large-scale models for the interpretation of changing regional patterns in biodiversity^3^. As a result, practitioners of ecosystem biomonitoring struggle to provide information that can easily be scaled up to interpret large-scale regional change^4^. This is a critical deficit, as ecosystems currently face significant threats arising from large-scale, pervasive environmental drivers such as climate change, which in turn create spatially and temporally diverse and co-acting stressors^5^.

Over the last decade, biodiversity science has experienced a genomics/bioinformatics revolution. The technique of DNA barcoding has supported the wider use of genetic information as a global biodiversity identification and discovery tool^6,7^. Several studies have advocated the use of DNA barcode sequences to identify bio-indicator species (e.g., macroinvertebrates) in the context of biomonitoring applications^8,9^. The use of DNA sequence information for specimen identification can significantly aid biomonitoring programs by increasing taxonomic resolution (which can provide robust species-level identification) in comparison to morphological analysis (which is often limited to genus- or family-level [order, or class-level] identification). However, this methodology still requires the sorting and separation of individual specimens from environmental samples obtained through collection methods such as benthic kick-net sampling. The samples obtained routinely contain hundreds to thousands of individual organisms, many of which are immature stages which cannot be reliably identified^10^.

Advances in high-throughput sequencing (HTS) technologies have enabled massively parallelized sequencing platforms with the capacity to obtain sequence information from biota in environmental samples without separating individual organisms^11,12^. Past research has demonstrated the utility of HTS in providing biodiversity data from environmental samples that have variously been called “metagenomics”, “environmental barcoding”, “environmental DNA” or “DNA metabarcoding”^12,13^. These approaches are either targeted towards specific organisms (e.g., pathogens, invasive species, or endangered species) or aim to characterize assemblages of biota. Biomonitoring applications fall mainly into the second category where assemblages are targeted for ecological analyses^3^. For example, macroinvertebrate larvae from benthos are considered standard bio-indicator taxa for aquatic ecosystem assessment. Previous work demonstrated the use of HTS in biodiversity analysis of benthic macroinvertebrates^14,15,16^ and its applicability to biomonitoring programs^3^. Various studies have contributed to this endeavor by demonstrating capabilities and limitations of HTS in aquatic biomonitoring^17,18,19,20^.

An important consideration in generating DNA information via HTS analysis for biomonitoring involves the choice of samples. A wide range of sample types including water, soil, benthic sediments, gut contents, passive biodiversity samplings (e.g., malaise traps) could be used as sources for DNA extraction and analysis^21^. Depending on the size of the target organisms, in some cases whole organisms might be present in the samples (e.g., larval samples in benthos). However, a sample may also harbor DNA in residual tissue or cells shed from organisms that may not be present as a whole. For example, early work on environmental DNA focused on detecting relatively large target species (e.g., invasive amphibian or fish species) from DNA obtained from water samples^22^. The idea of analyzing DNA obtained from water has been proposed for biodiversity assessment in and around water bodies or rivers including^23^ and specifically for bioindicator species^24^. However, because benthos harbors microhabitats for bio-indicator species development and growth, it has been the main source of biodiversity samples for biomonitoring applications^1^. In order to evaluate the suitability of water as a source for biodiversity information of bio-indicator taxa, it is important to assess whether DNA obtained from water samples alone provides sufficient coverage of benthic bio-indicator taxa commonly used in aquatic biomonitoring.

Here, we compare benthic and water samples collected in parallel from the same wetland ponds as sources of DNA for environmental DNA (eDNA) metabarcoding analysis. Specifically, we assess whether patterns of biodiversity illuminated through DNA analysis of benthos is reflected through DNA analysis of water samples. The study system involves two adjacent deltas in northern Alberta, Canada within Wood Buffalo National Park. By comparing patterns of sequence data from operational taxonomic units (OTUs) and multiple taxonomic levels (species, genus, family, and order), we explore differences between biodiversity data (i.e. taxonomic list information) from DNA extracted from water samples as compared to DNA extracted from co-located benthic samples.

## Methods

### Field sampling

Eight open-water wetland sites within the Peace-Athabasca delta complex were sampled in August 2011. All sites were located within Wood Buffalo National Park in Alberta, Canada. Full collection data are supplied in the Supplementary Material. Three replicate samples of the benthic aquatic invertebrate community (hereafter designated as ‘benthos’) were taken from the edge of the emergent vegetation zone into the submerged vegetation zone at each site. Replicated, paired samples were located approximately 100 metres apart. Samples were collected using a standard Canadian Aquatic Biomonitoring Network (CABIN) kick net with a 400*μ*m mesh net and attached collecting cup attached to a pole and net frame. Effort was standardized at two minutes per sample. Sampling was conducted by moving the net up and down through the vegetation in a sinusoidal pattern while maintaining constant forward motion. If the net became impeded by dislodged vegetation, sampling was paused so extraneous vegetation could be removed. Sampling typically resulted in a large amount of vegetation within the net. After sampling this vegetation was vigorously rinsed to dislodge attached organisms, and visually inspected to remove remaining individuals before discarding. The remaining material was removed from the net and placed in a white 1L polyethylene sample jar filled no more than half full. The net and collecting cup were rinsed and inspected to remove any remaining invertebrates. Samples were preserved in 95% ethanol in the field, and placed on ice in a cooler for transport to the field base. Here they were transferred to a freezer at – 20°C before shipment. A sterile net was used to collect samples at each site and field crew wore clean nitrile gloves to collect and handle samples in the field and laboratory, thereby minimizing the risk of DNA contamination between sites.

Three 1L water samples for subsequent DNA extraction were collected directly into sterile DNA/RNA free 1L polyethylene sample jars. Water samples were collected at the same locations as the benthos samples, immediately prior to benthic sampling to avoid disturbance, resulting in the resuspension of DNA from the benthos into the water column. Water samples were placed on ice prior to being transported to the lab.

### Water sample filtering and benthos homogenization

Under a positive pressure sterile hood, 1L water samples were filtered with 0.22 *μ*m filter (Mobio Laboratories). After water filtration, total DNA was extracted from the entire filter using Power water DNA extraction kit (MoBio Laboratories) and eluted in 100 *μ*l of molecular biology grade water, according to the manufacturer instructions. DNA samples were kept frozen at −20 C until further PCR amplification and sequencing. DNA extraction negative control was performed in parallel to ensure the sterility of the DNA extraction process.

For benthos samples, after removal of the EtOH^15^, a crude homogenate was produced by blending the component of each sample using a standard blender that had been previously decontaminated and sterilized using ELIMINase™ followed by a rinse with deionized water and UV treatment for 30 min. A representative sample of this homogenate was transferred to 50 mL Falcon tubes and centrifuged at 1000 rpm for 5 minutes to pellet the tissue. After discarding the supernatant, the pellets were dried at 70°C, until the ethanol was fully evaporated. Once dry, the homogenate pellets were combined into a single tube and stored at −20°C.

Using a sterile spatula, ~300 mg dry weight of homogenate was subsampled into 3 MP matrix tubes containing ceramic and silica gel beads. The remaining dry mass was stored in the Falcon tubes at −20°C as a voucher.

DNA was extracted using a NucleoSpin tissue extraction kit (Macherey-Nagel) with a minor modification of the kit protocol: the crude homogenate was first lysed with 720 *μ*L T1 buffer and then further homogenized using a MP FastPrep tissue homogenizer for 40 s at 6 m/s. Following this homogenization step, the tubes were spun down in a microcentrifuge and 100 *μ*L of proteinase K was added to each. After vortexing, the tubes were incubated at 56°C for 24 hr. Once the incubation was completed, the tubes of digest were centrifuged for 1 min at 10,000 g and 200 *μ*L of supernatant was transferred to each of three sterile microfuge tubes per tube of digest. The lysate was loaded to a spin column filter and centrifuged at 11,000 g for 1 min. The columns were washed twice and dried according to the manufacturer’s protocol. The dried columns were then transferred into clean 1.5 mL tubes. DNA was eluted from the filters with 30 *μ*L of warmed molecular biology grade water. DNA extraction negative control was performed in parallel to ensure the sterility of the DNA extraction process.

Purity and concentration of DNA for each site was checked using a NanoDrop spectrophotometer and recorded. Samples were kept at −20°C for further PCR and sequencing.

### Amplicon library preparation for HTS

Two fragments within the standard COI DNA barcode region were amplified with two primer sets (A_F/D_R [~250 bp] called AD and B_F/E_R called BE [~330 bp])^15,21^ using a two-step PCR amplification regime. The first PCR used *COI* specific primers and the second PCR involved Illumina-tailed primers. The PCR reactions were assembled in 25 *μ*l volumes. Each reaction contained 2 *μ*l DNA template, 17.5 *μ*l molecular biology grade water, 2.5 *μ*l 10× reaction buffer (200 mM Tris–HCl, 500 mM KCl, pH 8.4), 1 *μ*l MgCl2 (50 mM), 0.5 *μ*l dNTPs mix (10 mM), 0.5 *μ*l forward primer (10 mM), 0.5 *μ*l reverse primer (10 mM), and 0.5 *μ*l Invitrogen’s Platinum Taq polymerase (5 U/*μ*l). The PCR conditions were initiated with heated lid at 95°C for 5 min, followed by a total of 30 cycles of 94°C for 40 s, 46°C (for both primer sets) for 1 min, and 72°C for 30 s, and a final extension at 72°C for 5 min, and hold at 4°C. Amplicons from each sample were purified using Qiagen’s MiniElute PCR purification columns and eluted in 30 *μ*l molecular biology grade water. The purified amplicons from the first PCR were used as templates in the second PCR with the same amplification condition used in the first PCR with the exception of using Illumina-tailed primers in a 30-cycle amplification regime. All PCRs were done using Eppendorf Mastercycler ep gradient S thermalcyclers and negative control reactions (no DNA template) were included in all experiments.

### High throughput sequencing

PCR products were visualized on a 1.5% agarose gel to check the amplification success. All generated amplicons plates were dual indexed and pooled into a single tube. The pooled library were purified by AMpure beads and quantified to be sequenced on a MiSeq flowcell using a V2 MiSeq sequencing kit (250 × 2; FC-131-1002 and MS-102-2003).

### Bioinformatic methods

Raw Illumina paired-end reads were processed using the SCVUC v2.3 pipeline available from https://github.com/EcoBiomics-Zoobiome/SCVUC_COI_metabarcode_pipeline. Briefly, raw reads were paired with SeqPrep ensuring a minimum Phred score of 20 and minimum overlap of at least 25 bp^25^. Primers were trimmed with CUTADAPT v1.18 ensuring a minimum trimmed fragment length of at least 150 bp, a minimum Phred score of 20 at the ends, and allowing a maximum of 3 N’s^26^. All primer-trimmed reads were concatenated for a global ESV analysis. Reads were dereplicated with VSEARCH v2.11.0 using the ‘derep_fulllength’ command and the ‘sizein’ and ‘sizeout’ options^27^. Denoising was performed using the unoise3 algorithm in USEARCH v10.0.240^28^. This method removes sequences with potential errors, PhiX carry-over from Illumina sequencing, putative chimeric sequences, and rare reads. Here we defined rare reads to be exact sequence variants (ESVs) containing only 1 or 2 reads (singletons and doubletons)^29^ (Callahan, McMurdie, & Holmes, 2017). An ESV x sample table was created with VSEARCH using the ‘usearch_global’ command, mapping reads to ESVs with 100% identity. ESVs were taxonomically assigned using the COI Classifier v3.2^30^.

### Data analysis

Most diversity analyses were conducted in Rstudio with the *vegan* package^31,32^. Read and ESV statistics for all taxa and for arthropods only were calculated in R. To assess whether sequencing depth was sufficient we plotted rarefaction curves using a modified vegan *‘rarecurve’* function. Before normalization, we assessed the recovery of ESVs from benthos compared with water samples and assessed the proportion of all ESVs that could be taxonomically assigned with high confidence. Taxonomic assignments were deemed to have high confidence if they had the following bootstrap support cutoffs: species >= 0.70 (95% correct), genus >= 0.30 (99% correct), family >= 0.20 (99% correct) as is recommended for 200 bp fragments^30^. An underlying assumption for nearly all taxonomic assignment methods is that the query taxa are present in the reference database, in which case 95-99% of the taxonomic assignments are expected to be correct using these bootstrap support cutoffs. Assignments to more inclusive ranks, ex. order, do not require a bootstrap support cutoff to ensure that 99% of assignments are correct.

To assess how diversity recovered from benthos and water samples may differ, we first normalized different library sizes by rarefying down to the 15^th^ percentile library size using the vegan ‘*rrarefy*’ function^33^. It is known that bias present at each major sample-processing step (DNA extraction, mixed template PCR, sequencing) can distort initial template to sequence ratios rendering ESV or OTU abundance data questionable^17,34,35,36^. Here we chose to transform our abundance matrix to a presence-absence matrix for all further analyses. We calculated ESV richness across different partitions of the data to compare differences across sites and collection methods (benthos or water samples). To check for significant differences we first checked for normality using visual methods (*ggdensity* and *ggqqplot* functions in R) and the Shapiro-Wilk test for normality^37^. Since our data was not normally distributed, we used a paired Wilcoxon test to test the null hypothesis that median richness across sites from benthic samples is greater than the median richness across sites from water samples^38^.

To assess the overall community structure detected from different collection methods, we used non-metric multi-dimensional scaling analysis on Sorensen dissimilarities (binary Bray-Curtis) using the vegan ‘*metaMDS*’ function. A scree plot was used to guide our choice of 3 dimensions for the analysis (not shown). A Shephard’s curve and goodness of fit calculations were calculated using the vegan ‘*stressplot*’ and ‘*goodness*’ functions. To assess the significance of groupings, we used the vegan *‘vegdist* function to create a Sorensen dissimilarity matrix, the ‘*betadisper*’ function to check for heterogeneous distribution of dissimilarities, and the ‘*adonis*’ function to do a permutational analysis of variance (PERMANOVA) to check for any significant interactions between groups (collection method, sample site). We calculated the Jaccard index to look at the overall similarity between water and benthos samples.

To assess the ability of traditional bioindicator taxa to distinguish among samples, we limited our dataset to ESVs assigned to the EPTO (Ephemeroptera, Plecoptera, Trichoptera, Odonata) insect orders. No significant beta dispersion was found within groups. We used PERMANOVA to test for significant interactions between groups and sources of variation such as collection method and river delta as described above. Sample replicates were pooled. We also visualized the frequency of ESVs detected from EPTO families using a heatmap generated using *geom_tile* (ggplot) in R.

## Results

A total of 48,799,721 x 2 Illumina paired-end reads were sequenced (Table S1). After bioinformatic processing, we retained a total of 16,841 ESVs (5,407,720 reads) that included about 11% of the original raw reads. Many reads were removed during the primer-trimming step from water samples for being too short (< 150 bp) after primer trimming. After taxonomic assignment, a total of 4,459 arthropoda ESVs (4,399,949 reads) were retained for data analysis (Table S2). 27% of all ESVs were assigned to arthropoda, accounting for 81% of reads in all ESVs.

Rarefaction curves that reach a plateau show that our sequencing depth was sufficient to capture the ESV diversity in our PCRs (Figure S1). Benthos samples generate more ESVs than water samples as shown in the rarefaction curves as well as by the median number of reads and ESVs recovered by each collection method (Figure S2). As expected, not all arthropoda ESVs could be taxonomically assigned with confidence (Figure S3). This is probably because local arthropods may not be represented in the underlying reference sequence database. As a result, most of our analyses are presented at the finest level of resolution using exact sequence variants.

### Analysis of sample biodiversity

Alpha diversity measures based on mean richness and beta diversity based on the Jaccard index among all samples show higher values for benthos compared to water at the ESV rank. The total richness for benthos is 1,588 and water is 658, with a benthos:water ratio of 2.4. The Jaccard index is 0.14 indicating that water and benthos samples are 14% similar. Examining the arthropod ESV richness for each sample site from benthos and water collections reinforces the general pattern of higher detected richness from benthos samples (Wilcoxon test, p-value < 0.05) (Figure 1).

**Figure 1.**
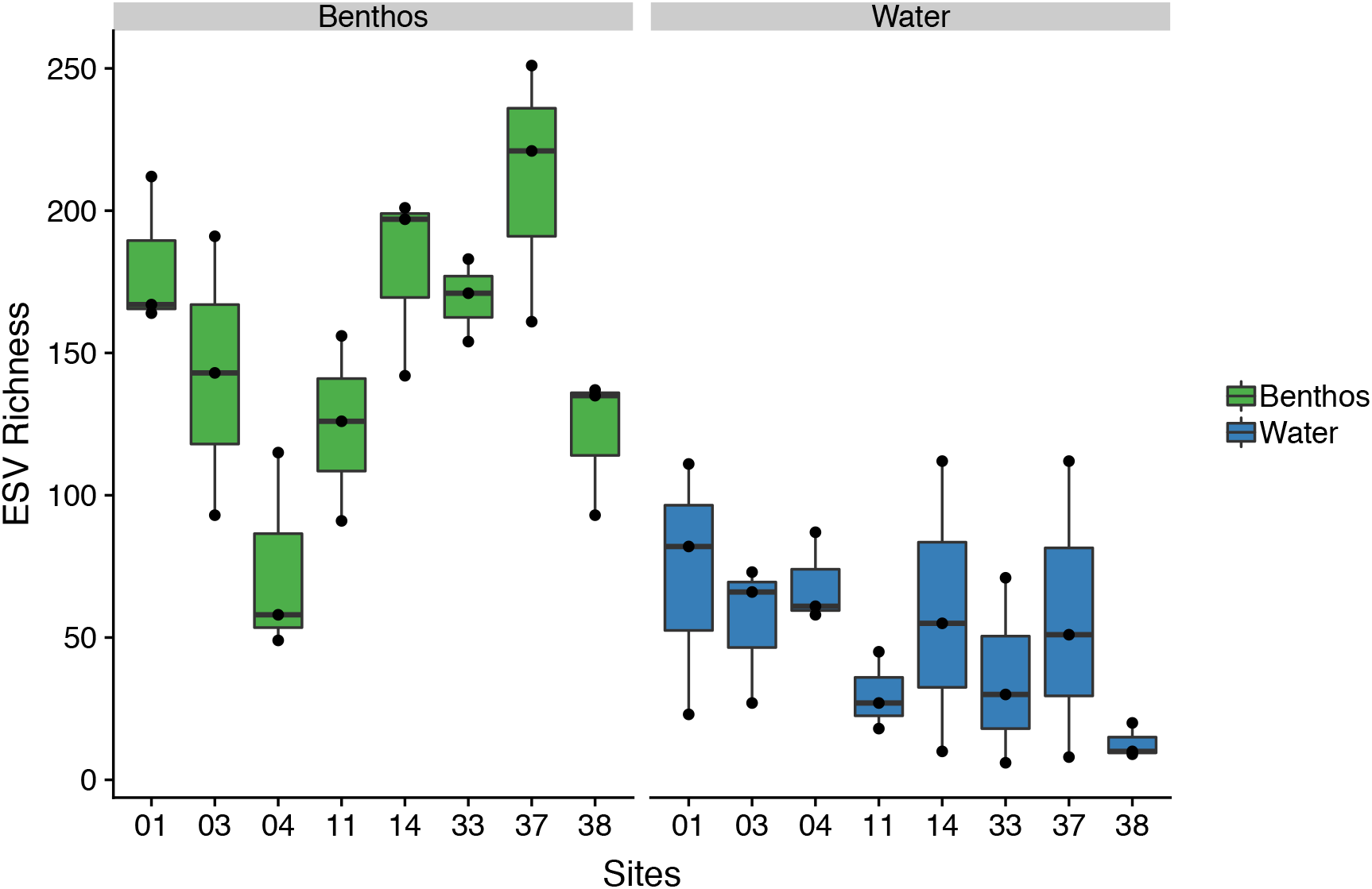
Median arthropod richness per site is higher in benthos samples than water samples. Results are based on normalized data.

We further illustrate how arthropod richness varies with collection method (benthos or water) by looking at the number of ESVs exclusively found from benthos samples, found both benthos and water samples, or exclusively found from water samples (Figure 2). For example, for sample 04B, 49% of ESVs are unique to benthos samples, 37% of ESVs are unique to water samples, and 14% of ESVs are shared. In fact, this sample contains the largest proportion of shared ESVs. When looking at more inclusive taxonomic ranks, more of the community is shared among benthos and water samples. When considering specific arthropod orders and genera, a greater diversity of sequence variants are detected from benthic samples even when the same higher-level taxa are also recovered from water samples (Figure 3). Some of the confidently identified arthropod genera represented by more than 100 sequence variants included: *Tanytarsus* (Diptera identified from benthos-B and water-W), *Aeshna* (Odonata, B only), *Leucorrhinia* (Odonata, B only), and *Scapholeberis* (Diplostraca, B + W).

**Figure 2.**
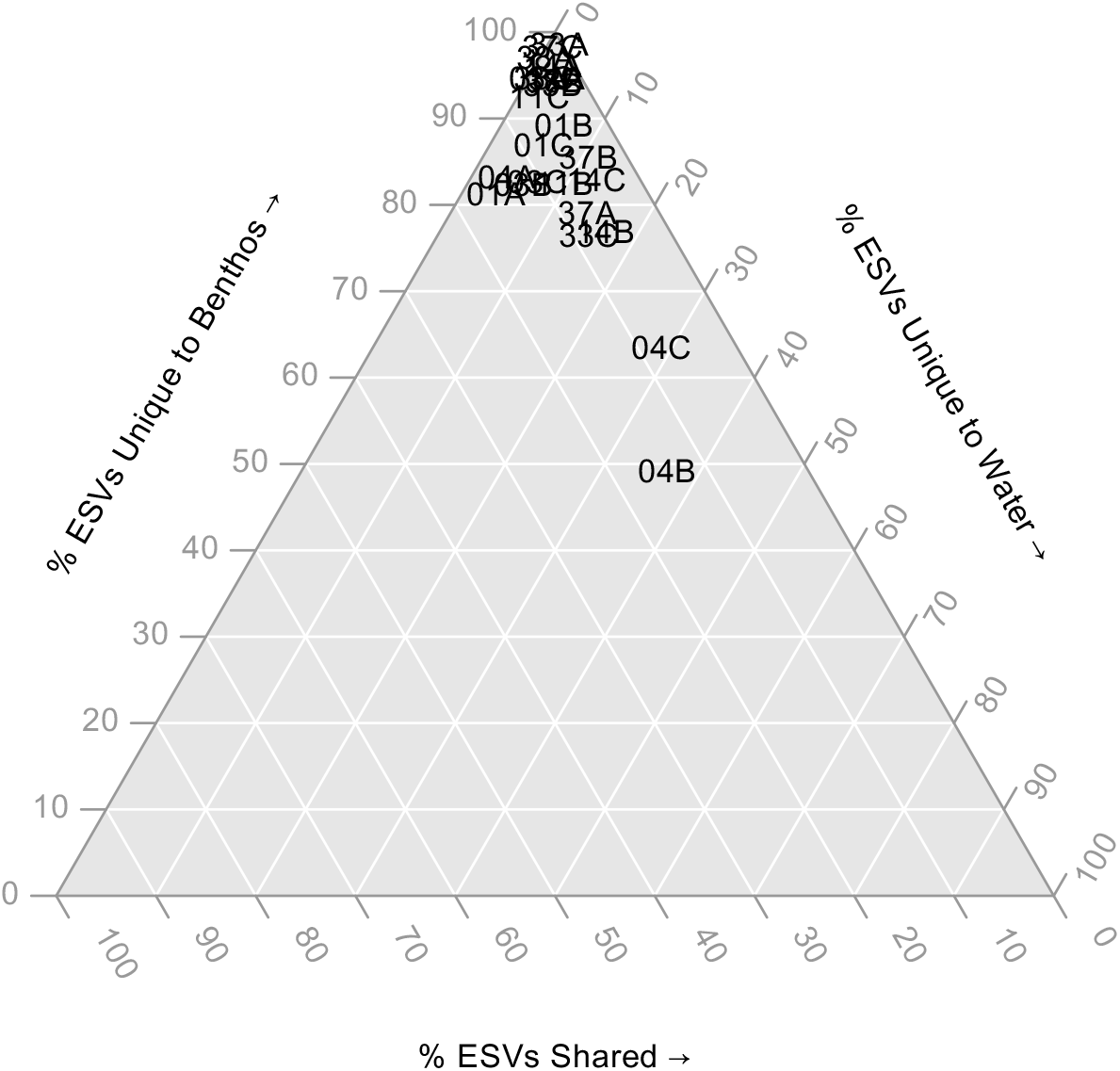
Few arthropod ESVs are shared among benthic and water samples. The ternary plot shows the proportion of ESVs unique to benthos samples, unique to water samples, or shared. Sample names are shown directly on the plot. Results are based on normalized data.

**Figure 3.**
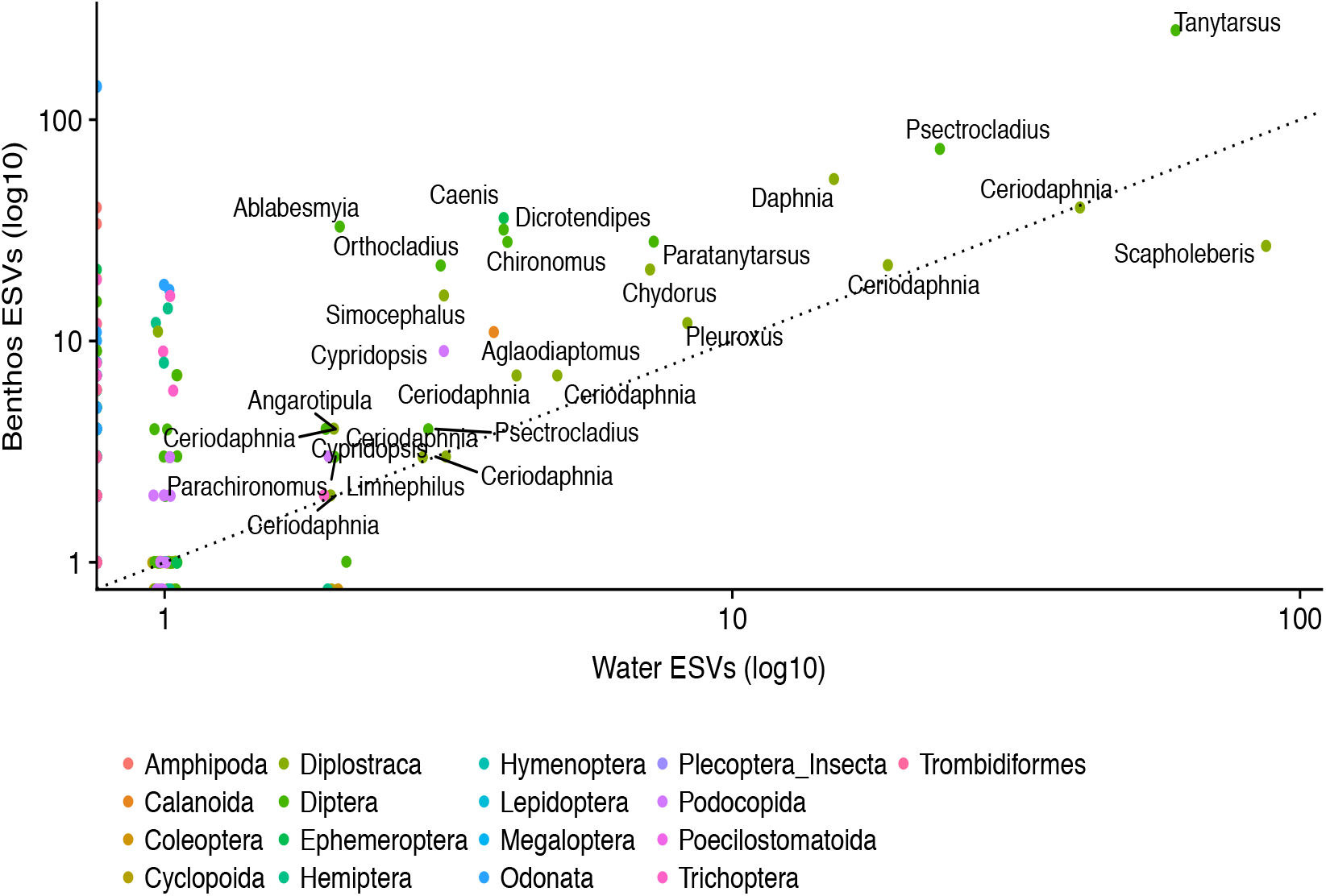
A greater diversity of arthropod sequence variants are detected from benthic samples. Each point represents a genus identified with high confidence and the number of benthic and water exact sequence variants (ESVs) with this taxonomic assignment. Only genera represented by at least 2 ESVs in both benthic and water samples are labelled in the plot for clarity. The points are color coded for the 17 arthropod orders detected in this study. A 1:1 correspondence line (dotted) is also shown. Points that fall above this line are represented by a greater number of ESVs from benthic samples. A log10 scale is shown on each axis to improve the spread of points with small values.

Samples from the same sites, but collected using different methods (benthos or water), clustered according to collection method intead of site (Figure 4). The ordination was a good fit to the observed Sorensen dissimilarities (NMDS, stress = 0.12, R^2^ = 0.91). Visually, samples cluster both by collection method and river delta. Although we did find significant beta dispersion among collection method, river, and site dissimilarities (ANOVA, p-value < 0.01), we had a balanced design so we used a PERMANOVA to check for any significant interactions between groups and none were found^39^. Collection site explained ~ 53% of the variance (p-value < 0.05), river delta explained ~ 10% of the variance (p-value = 0.001), and collection method explained ~ 9% of the variance in beta diversity (p-value = 0.001). Thus, even though richness measures are highly sensitive to choice of collection method, beta diversity is robust with samples clearly clustering by river delta regardless of whether benthos or water samples are analyzed (p-value = 0.001; Figure S4).

**Figure 4.**
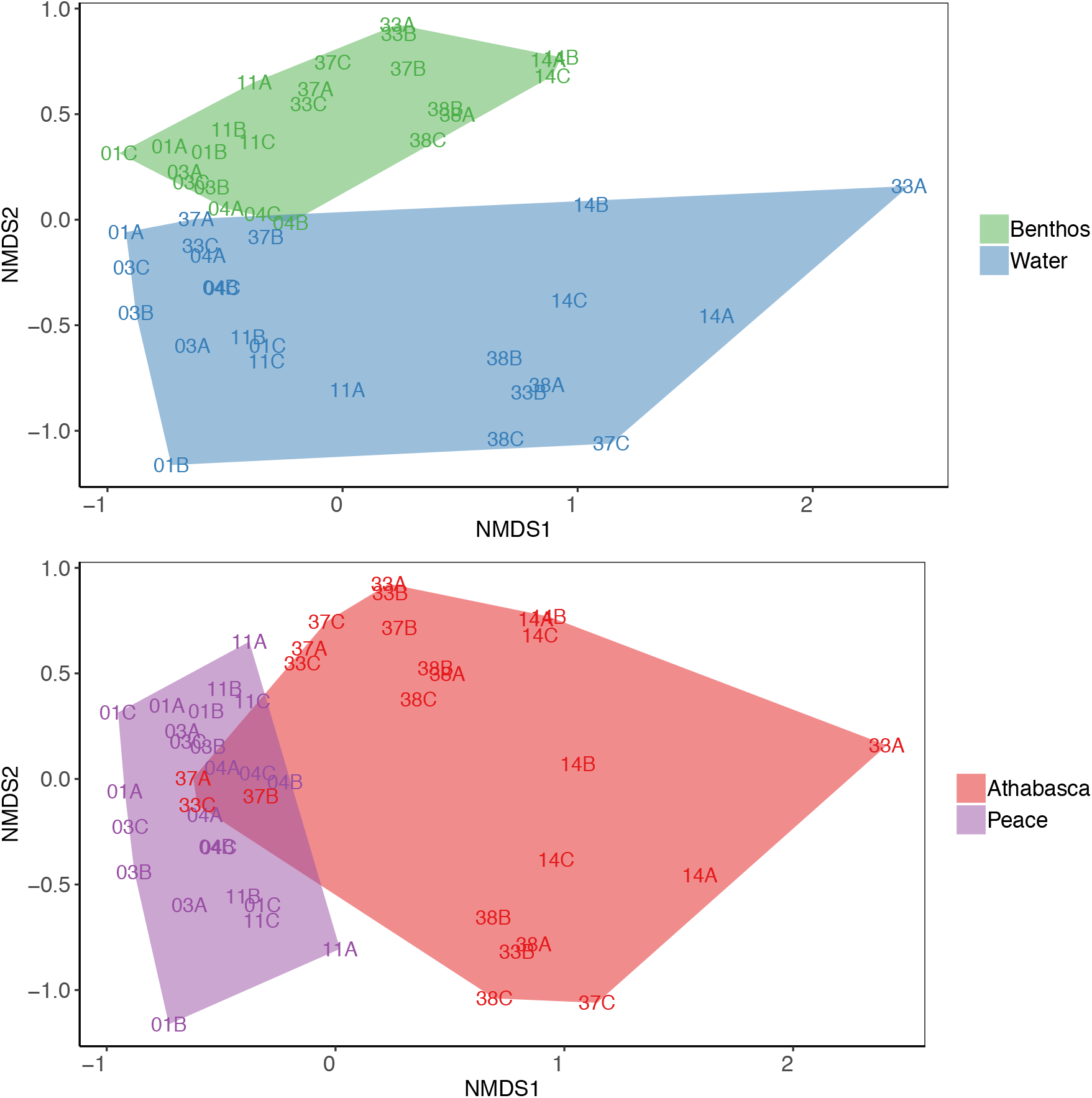
Samples cluster by collection method and river delta. The NMDS is based on rarefied data and Sorensen dissimilarities based on presence-absence data. The first plot shows sites clustered by collection method, benthos or water. The second plot shows sites clutered by river delta, Athabasca River or Peace River delta.

### Analysis of key bioindicator groups

Given the importance of aquatic insects as bioindicator species in standard biomonitoring programs, and to specifically address whether water samples could be used in lieu of benthos for biomonitoring applications, we closely examined the results obtained for four insect orders of biomonitoring importance. Based on the detection of EPTO ESVs, collection method (benthos or water) accounts for 13% of the variation in ordination distances (PERMANOVA, p-value=0.011; Table S3). Overall, these differences stem from variation in the distribution of ESVs detected from 76 observed EPTO families (Figure 5). While the total number of ESVs and EPTO families varied from site to site, there is a dramatic shift in the composition detected from benthos and water. For example, in site 1, 888 ESVs from 40 EPTO families were detected from the benthos sample, while only 133 ESVs from 9 EPTO families were observed from the water sample, despite being taken at the exact same location and time. Within each collection method, river delta explains 11% of the variation (PERMANOVA, p-value=0.031). This means is that despite differences in the community composition detected from benthos and water, EPTO ESVs can still be used to separate samples from two river deltas.

**Figure 5.**
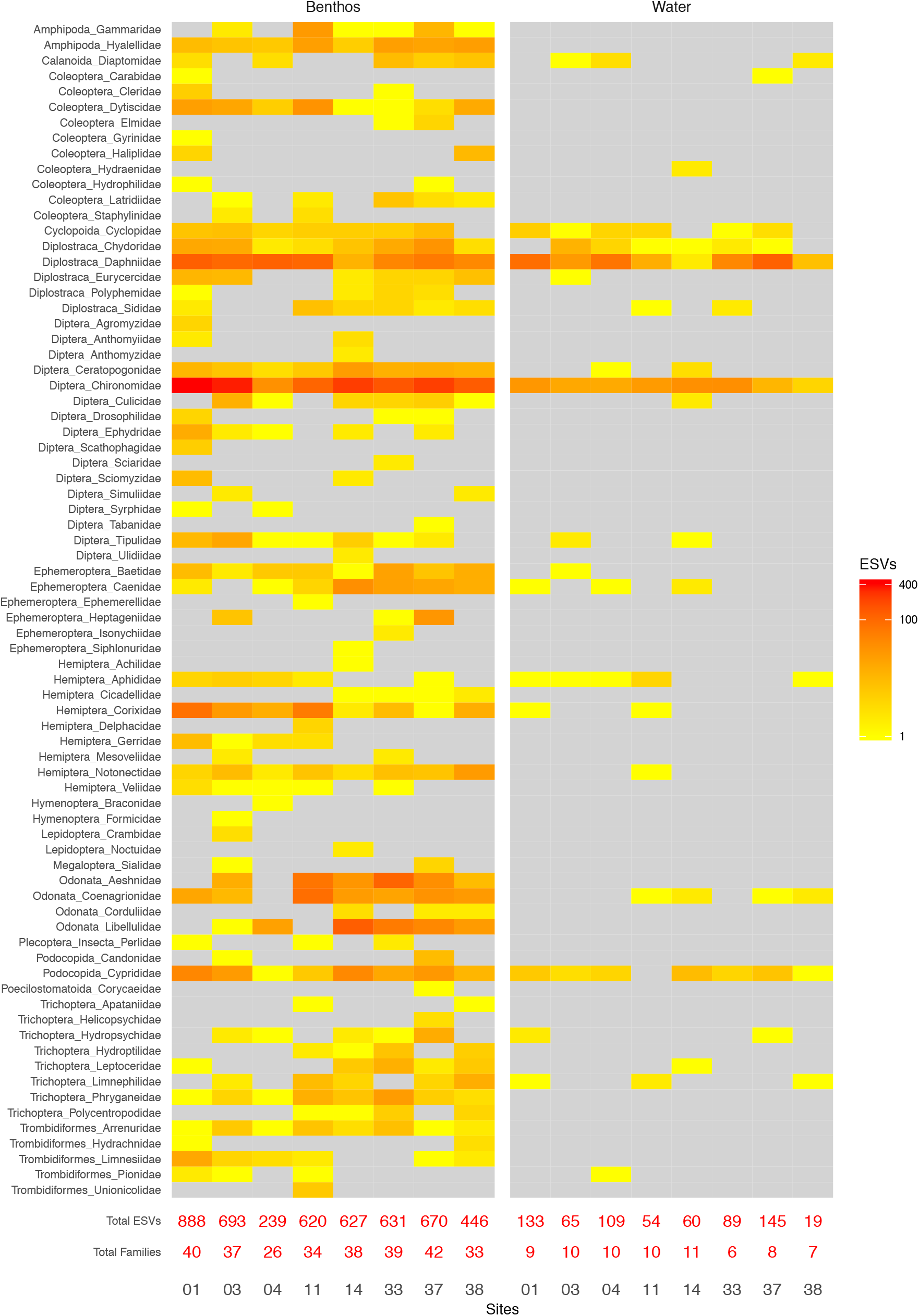
More Ephemeroptera, Plecoptera, Trichoptera, and Odonata family ESVs are detected from benthos compared with water samples. Each cell shows ESV richness colored according to the legend. Grey cells indicate zero ESVs. Only ESVs taxonomically assigned to families with high confidence (bootstrap support >= 0.20) are included. Based on normalized data.

## Discussion

Biodiversity information forms the basis of a vast array of ecological and evolutionary investigations. Given that biodiversity information for bioindicator groups, such as aquatic insects, is the main source of biological data for various environmental impact assessment and monitoring programs, it is vital for these data to provide a consistent and accurate representation of existing taxon richness^40^. Methods based on bulk sampling of environmental material (i.e. water) for identification of either single species^41^ or communities^42^, has been proposed as a simplified biomonitoring tool^23,24,43^. However, our analysis shows that water eDNA fails to provide a rich representation of the community structure in aquatic ecosystems. Our unique sampling design allowed us to undertake a direct comparison as we were able to collect samples from benthos and water in parallel across a range of sites. These wetland sites consisted of small ponds with minimal or no flow, minimizing the chance of stream flow as a factor impacting the availability of eDNA in a given water sample.

Our analysis of taxon richness in benthos versus water illuminates the need for caution when interpreting data captured from water as an estimate of total richness in a system. In some cases, we saw several-fold decreases in richness in water versus benthos. Although a comprehensive analysis of taxon richness should not rely solely on numbers, this reduction in taxa detected indicates the inadequacy of water for solely detecting existing aquatic invertebrate communities. In comparison, a recent study suggested that eDNA metabarcoding in flowing systems recovers higher levels of richness than bulk benthos samples^44^. However, our study design allowed a direct comparison between water and benthos for both EPTO and general richness without the influence of flow, meaning this was a true assessment of local community assemblages, represented by each sample type. eDNA metabarcoding in flowing systems can therefore result in the additional detection of upstream communities^44^, reflected in the greater number of taxa detected, but does not reflect the existing biodiversity at the local scale.

An important consideration when deciding effective biomonitoring methods should be the ecology of target biodiversity units. Factors including life cycle and habitat preference (i.e. benthic or water column) is likely to influence the rate of detection in different sampling approaches^45,46,47^. We have demonstrated in this study that whilst some ESVs are shared between both benthos and water, there is a sampling bias as to the associations of taxa, particularly EPTO, with different sample sources, which was also observed in a recent comparative study with running water^44^. The association of specific taxa with benthos enables communities to be assessed spatially, across different habitat types^14,48^. One of the major limitations of attempting to determine presence/absence of taxa in water is the uncertainty of the original DNA source. As samples are often collected at single fixed locations, taxa recovered in water can vary depending on when and where DNA was released into the aqueous environment in addition to other factors including flow rate^23^. This makes scaling up results from water challenging^49^. Conversely, benthos samples enable a real-time assessment of biodiversity originating from a known locality, which has implications for fine-scale environmental assessments^14^.

Environmental factors including hydrolysis drive DNA degradation in aqueous substrates, which can negatively influence detectability of DNA in water^50^. This confounding factor could account for some of the reduction in biodiversity observed between benthos and water^51^. For water sampling to improve species coverage and gain a comparable number of observations, a considerable increase in replicates and sequencing depth is required^52,53^. Earlier research has shown that increasing the volume of water up to 2 L does not seem to be a factor in additional taxonomic coverage^24^, however increasing the number of both biological and technical replicates can increase the number of taxa detected^52,53,55^. We used triplicate sampling for each site and compared EPTO taxa between sites and two rivers, separately. None of these comparisons provided support for the use of water eDNA in place of benthos. We found that benthos replicates clustered closer with less variation in ESV abundance in comparison with water, which suggests that three replicates is sufficient for consistent species detection with benthos and water is less consistent at representing community structure. In addition, using highly degenerate primers can increase the total biodiversity detected using eDNA metabarcoding^44^. However, with highly degenerated primers, there is an increase likelihood of amplifying non-target regions^56^, in comparison to primers with lower degeneracy such as those used in this study. Additionally, employing highly degenerate primers in biomonitoring studies lead to overrepresentation of some taxa (e.g. non-metazoan), which further distances such metabarcoding studies from current stream ecosystem assessment methods^44,57^. Attempting to improve taxonomic coverage of water by increasing numbers of samples collected, sequencing depth and utilising highly degenerate primers, adds considerable costs, both financial, in terms of effort and comparability, without the guarantee of representative levels of biodiversity identification.

## Conclusions

It is apparent that in data generated from our comparative study, employing water column samples as a surrogate for true benthic samples is not supported, as benthos DNA does not appear to be well represented in the overlying water in these static-water wetland systems at detectable levels. Benthic samples are a superior source of biomonitoring DNA when compared to water in terms of providing reproducible taxon richness information at a variety of spatial scales. Choice of sampling method is a critical factor in determining the taxa detected for biomonitoring assessment and we believe that a comprehensive assessment of total biodiversity should include multiple sampling methods to ensure that representative DNA from all target organisms can be captured.

## Acknowledgements

T.M. Porter was suppored by the Government of Canada through the Genomics Research and Development Initiative (GRDI), Ecobiomics project. We acknowledge field support from Parks Canada (Jeff Shatford, Ronnie & David Campbell) and Environment and Climate Change Canada (Daryl Halliwell) for the process of sample collection. This study is funded by the government of Canada through Genome Canada and Ontario Genomics.

## Author contributions and competing interests

M.H. designed the overall study, contributed to genomics and bioinformatics analyses, and wrote the manuscript; T.M.P. conducted bioinformatic processing, helped analyze data, and helped to write the manuscript; S.S. conducted molecular biology and genomics analyses; C.V.R. assisted with interpreting data and helped to write the manuscript. D.J.B. planned and organised the field study, collected the samples and helped to write the manuscript; M.W. conducted molecular biology and genomics analyses. The authors declare no competeing interests.

## Data availability

Raw sequences will be deposited to the NCBI SRA on acceptance. A FASTA file of ESV sequences are available in the Supplmentary Material. The bioinformatic pipeline and scripts used to create figures are available on GitHub at https://github.com/terrimporter/CO1Classifier.

