## Supplementary Information for "Watered-down biodiversity? A comparison of metabarcoding results from DNA extracted from matched water and bulk tissue biomonitoring samples"

**Table S1. Summary of reads and ESVs in all taxa**

| | AD | | BE | | |
|---|---|---|---|---|---|
| | **Benthos** | **Water** | **Benthos** | **Water** | **Total** |
| Raw | N/A | | N/A | | 48,799,721 x 2 |
| Paired | N/A | | N/A | | 42,317,963 |
| Primer Trimmed | 4,377,830 | 345,014 | 4,001,465 | 1,561,360 | 8,724,309 |
| ESVs | 2,581 | 1,099 | 4,831 | 9,571 | 16,841 |
| Reads in ESVs | 2,614,558 | 238,697 | 1,774,386 | 780,079 | 5,407,720 |
| Proportion of raw reads in ESVs (%) | 5.4 | 0.5 | 3.6 | 1.6 | 11.1 |

**Table S2. Summary of reads and ESVs assigned to the Arthropoda**

| | AD | | BE | | |
|---|---|---|---|---|---|
| | **Benthos** | **Water** | **Benthos** | **Water** | **Total** |
| ESVs | 1,735 | 280 | 2,398 | 491 | 4,459 |
| Reads in ESVs | 2,541,062 | 174,605 | 1,554,853 | 129,429 | 4,399,949 |
| Proportion of raw reads in ESVs (%) | 5.2 | 0.4 | 3.2 | 0.3 | 9.0 |
| Proportion of all ESVs that are Arthropoda | 67.2 | 25.5 | 49.6 | 5.1 | 26.5 |
| Proportion of all reads in ESVs that are Arthropoda | 97.2 | 73.1 | 87.6 | 16.6 | 81.4 |

**Table S3. EPTO ESVs can be used to separate rivers using either benthos or water samples.** Sample replicates were pooled. No significant beta dispersion was detected within groups (collection method, river). No significant interaction between groups was detected (collection method, river). Summary of PERMANOVA results based on a Sorensen dissimilarity matrix of EPTO ESVs. Significant p-values are in bold. Based on normalized data.

| Source of variation | Df | MS | F | $R^2$ | P |
|---|---|---|---|---|---|
| A) Interaction between groups | | | | | |
| Collection method | 1 | 0.80 | 2.09 | 0.13 | **0.002** |
| River | 1 | 0.66 | 1.73 | 0.11 | **0.021** |
| Collection method : River | 1 | 0.44 | 1.16 | 0.07 | 0.282 |
| Residuals | 11 | 0.38 | | 0.69 | |
| Total | 14 | | | 1.00 | |
| B) Variation due to collection method | | | | | |
| Collection method | 1 | 0.80 | 1.95 | 0.13 | **0.011** |
| Residuals | 13 | 0.41 | | 0.87 | |
| Total | 14 | | | 1.00 | |
| C) Within each collection method, variation due to river | | | | | |
| River, Stratum = Collection Method | 1 | 0.65 | 1.56 | 0.11 | **0.031** |
| Residuals | 13 | 0.41 | | | |
| Total | 14 | | | | |

Df = Degrees of freedom; MS = MeanSqs; F = F.Model; P = P-value

**Figure S1. Rarefaction curves are saturated.** Benthos samples from each site are shown in green and water samples are shown in blue. The vertical line shows the number of reads that would be included after normalizing library size down to the 15$^{th}$ percentile (reads = 2,099).

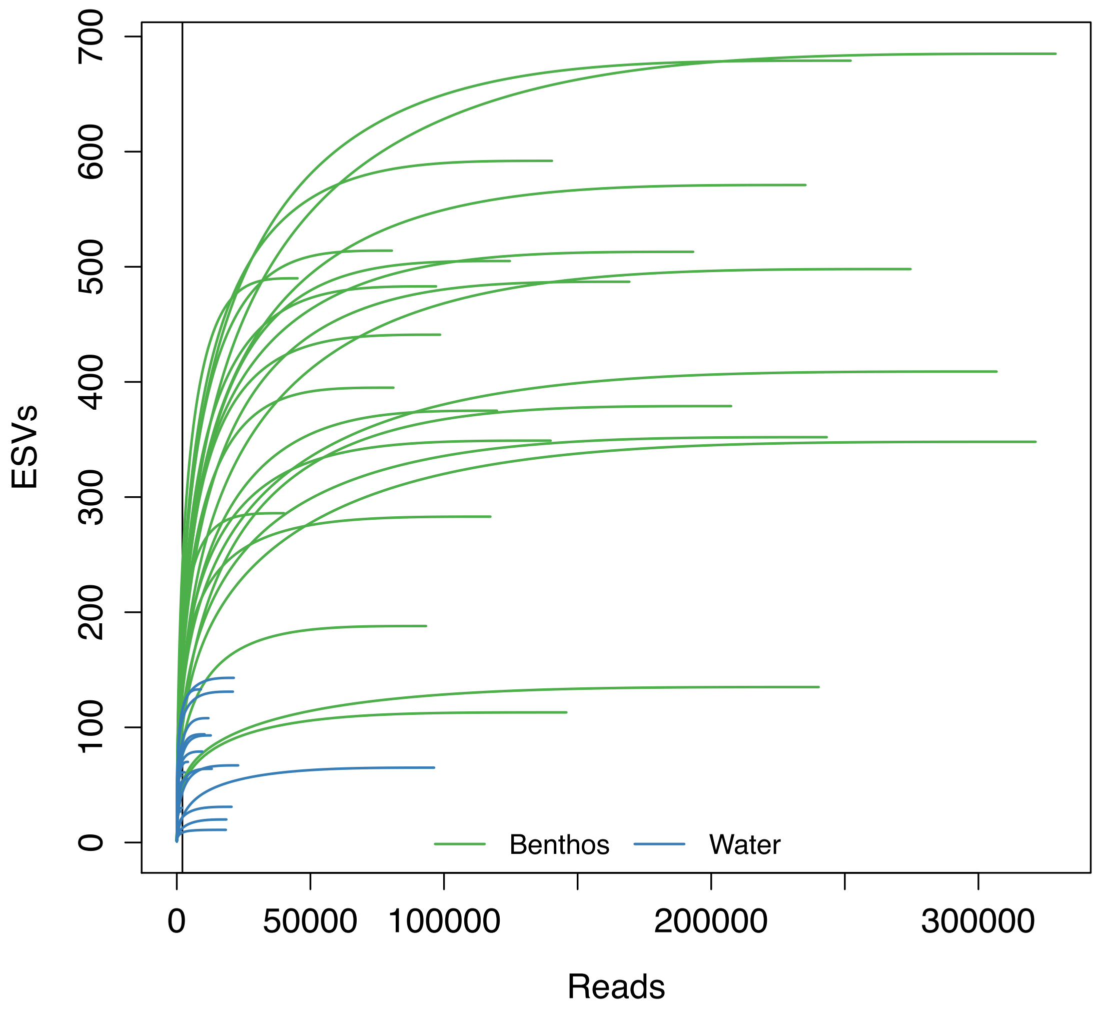

**Figure S2. The recovery of reads and ESVs from benthos is much greater than that from water.** Results summarized before normalization.

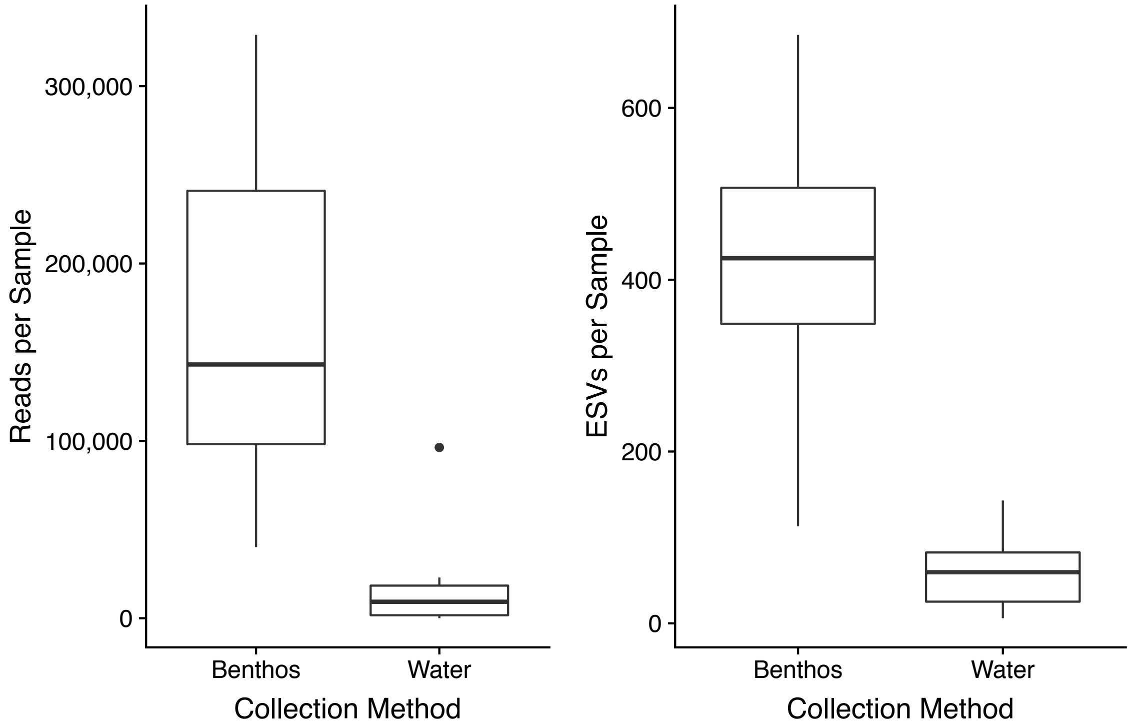


**Figure S3. Only some Arthropod ESVs could be assigned with high confidence.** Results summarized before normalization.

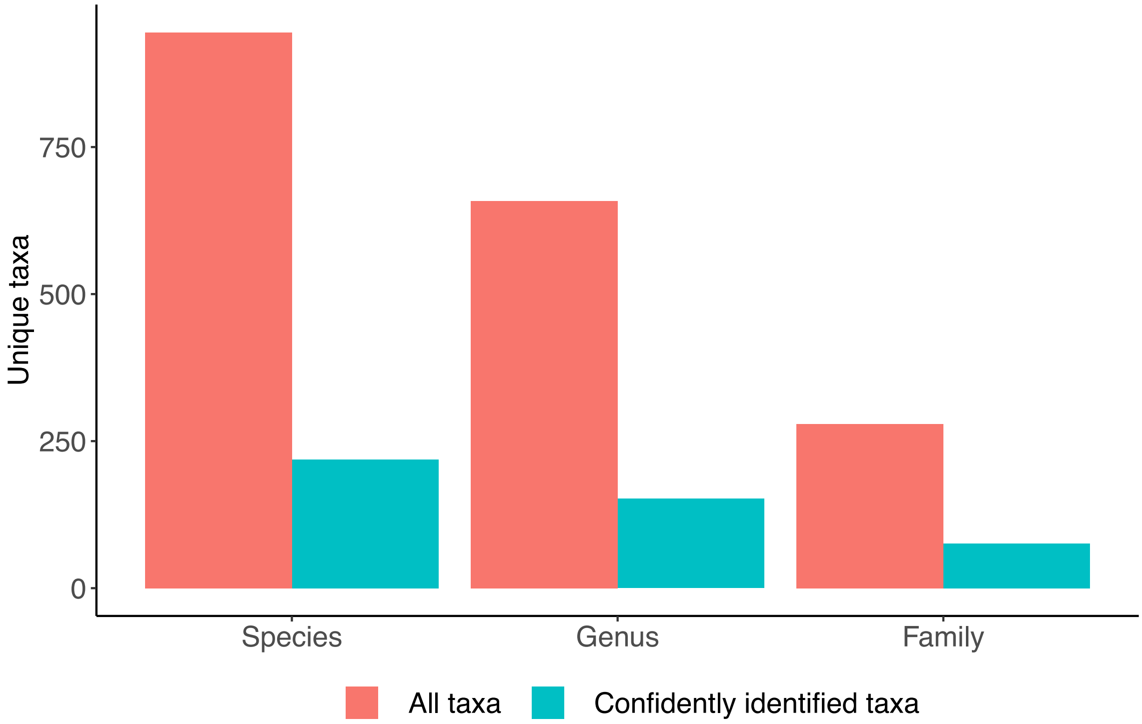

**Figure S4. River sites are separated whether Arthropod ESVs are detected from benthos or water samples.** NMDS ordination distances well-represent observed Sorensen dissimilarities (Benthos stress = 0.08, $R^2$ = 0.95; Water stress = 0.09, $R^2$ = 0.95). PERMANOVA shows that river groupings are significant and explain 14-19% of the variation in beta diversity (Benthos $R^2$ = 0.19, p-value = 0.001; Water $R^2$ = 0.14, p-value = 0.001). Results based on normalized data.

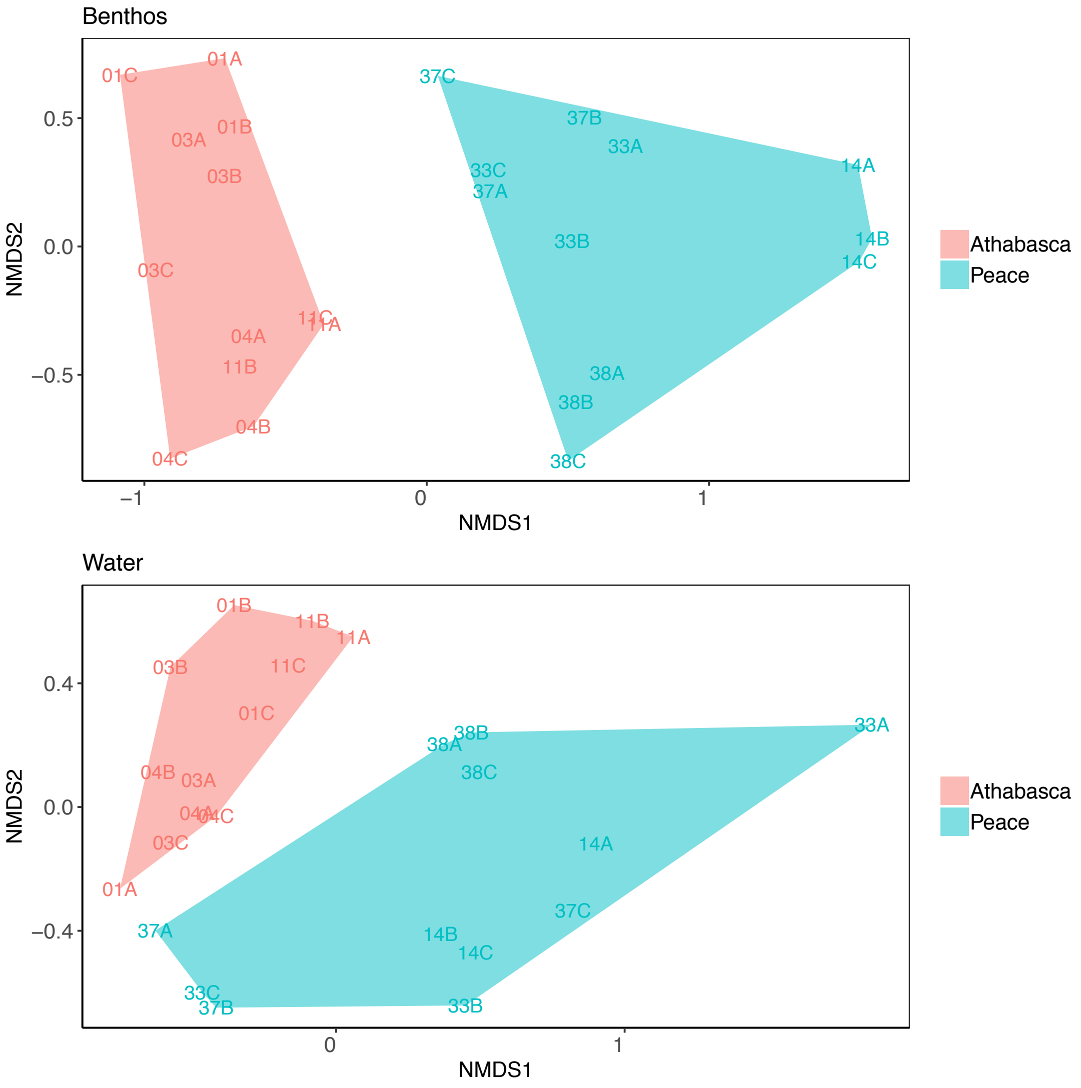
